# OsIRO3 negatively regulates Fe homeostasis by repressing the expression of *OsIRO2*

**DOI:** 10.1101/2022.02.14.480324

**Authors:** Chenyang Li, Yang Li, Peng Xu, Gang Liang

**Author notes:** Correspondence: Gang Liang.

## Abstract

Iron (Fe) is crucial for crop productivity and quality. However, Fe deficiency is prevalent worldwide, especially in alkaline soil. Plants have evolved sophisticated mechanisms to withstand Fe deficiency conditions. *Oryza sativa* IRON-RELATED BHLH TRANSCRIPTION FACTOR 3 (OsIRO3/OsbHLH63) has been identified as a negative regulator of Fe deficiency response signaling, however, the underlying mechanism remains unclear. In the present study, we constructed two *iro3* mutants which generated leaves with necrotic lesions under Fe deficient conditions. Loss-of-function of *OsIRO3* caused upregulation of Fe deficiency-associated genes in the root under Fe deficient conditions. Fe concentration measurement showed that the *iro3* mutants had increased shoot Fe concentration only under Fe deficient conditions. Further analysis revealed that OsIRO3 directly regulated the expression of *IRON-RELATED BHLH TRANSCRIPTION FACTOR 2* (*OsIRO2*) which encodes a positive regulator of Fe uptake system. Protein interaction tests indicated that OsIRO3 interacted with OsPRI1 and OsPRI2. Further investigation demonstrated that OsIRO3 repressed the transactivation of OsPRI1 and OsPRI2 towards *OsIRO2*. OsIRO3 contains an EAR motif which recruits the TOPLESS/TOPLESS-RELATED (OsTPL/OsTPRs) corepressors. Mutation of the EAR motif attenuated the repression ability of OsIRO3. This work sheds light on the molecular mechanism by which OsIRO3 modulates Fe homeostasis in rice.

## INTRODUCTION

Iron (Fe) is one of the indispensable micronutrients for plant growth and development, which is involved in many physiological and biochemical reactions such as photosynthesis, mitochondrial respiration, hormone biosynthesis, nitrogen fixation and so on (Hänsch and Mendel, 2009; Balk and Schaedler, 2014). Although Fe is abundant on earth, its availability is limited due to the low solubility at alkaline pH (Mori, 1999). Calcareous soil accounts for about one third of the world’s cultivated soil, making Fe deficiency a very common phenomenon (Guerinot and Yi, 1994). Fe deficiency often leads to interveinal chlorosis of leaves, as well as greatly affecting the yield and nutritional quality of crops (Briat et al., 2015). Reactive oxygen radicals produced by excess Fe are toxic to plant cells (Valko et al., 2005). Therefore, the Fe concentration in plant cells needs to be regulated strictly.

To cope with Fe deficiency, plants have developed complicated molecular mechanisms for Fe uptake, translocation, and storage to meet Fe demand. Plants have evolved different strategies to absorb Fe (Römheld and Marschner, 1986). Gramineous plants employ a chelation strategy to acquire Fe. They excrete mugineic acid family phytosiderophores (MAs) to chelate Fe^3+^ to form MA-Fe^3+^ complex which is translocated into roots by YS/YSL transporters. In rice, the synthesis of MAs is mediated by a series of enzymes, including S-adenosylmethionine synthetase (SAMS), nicotianamine synthase (NAS), nicotianamine aminotransferase (NAAT), and deoxymugineic acid synthase (DMAS) (Shojima et al., 1990; Mori 1999; Bashir et al., 2017). The efflux of MAs from roots counts on TRANSPORTER OF MAs 1 (OsTOM1) (Nozoye et al., 2011) and the influx of Fe^3+^-MA to roots involves YELLOW STRIP LIKE 15 (OsYSL15) (Inoue et al., 2009; Lee et al., 2009). In addition to the chelation strategy, rice plants also directly acquire Fe^2+^ by the Fe^2+^ transporter OsIRT1 (Ishimaru et al., 2006).

The Fe deficiency response is under the control of a series of transcription factors which constitute a complex regulatory network. IRON-RELATED BHLH TRANSCRIPTION FACTOR 2 (OsIRO2) is a key positive transcription factor of Fe homeostasis, which positively modulates the expression of chelation strategy associated genes including *OsNAS1*, *OsNAS2*, *OsNAAT1*, *OsDMAS1*, *OsTOM1*, and *OsYSL15* (Ogo et al., 2007; Liang et al., 2020; Wang et al., 2020). *Oryza sativa* FER-LIKE FE DEFICIENCY-INDUCED TRANSCRIPTION FACTOR (OsFIT)/OsbHLH156 was identified as an interacting partner of OsIRO2. OsIRO2 protein mainly localizes to the cytoplasm, and OsFIT can facilitate the nuclear accumulation of OsIRO2 under Fe limited conditions (Liang et al., 2020; Wang et al., 2020). OsFIT and OsIRO2 interdependently regulate the expression of chelation strategy associated genes (Liang et al., 2020). *OsIRO2* is inducible by Fe deficiency both in the root and shoot (Ogo et al., 2007), and its upregulation is dependent on *Oryza sativa* POSITIVE REGULATOR OF IRON HOMEOSTASIS (OsPRI) proteins, OsPRI1 (OsbHLH60), OsPRI2 (OsbHLH58), and OsPRI3 (OsbHLH59) (Zhang et al., 2017, 2020; Kobayashi et al., 2019). *Oryza sativa* HEMERYTHRIN MOTIF-CONTAINING REALLY INTERESTING NEW GENE AND ZINC-FINGER PROTEIN1 (OsHRZ1) and OsHRZ2 are two potential Fe sensors, which negatively regulate the expression of Fe deficiency inducible genes (Kobayashi et al., 2013). OsHRZ1 possesses a RING domain responsible for its E3 ligase activity. Recently, it is established that OsHRZ1 interacts with OsPRI1/2/3 and promotes the degradation of the latter (Zhang et al., 2017, 2020).

OsIRO3 was identified as a nuclear-localized negative regulator of Fe homeostasis (Zheng et al., 2010). Similar to OsIRO2, OsIRO3 is also inducible under Fe deficient conditions and directly regulated by OsPRI1/2/3 (Zhang et al., 2017, 2020; Kobayashi et al., 2019). Overexpression of *OsIRO3* causes leaf chlorosis, reduced shoot Fe concentration, and downregulation of Fe deficiency inducible genes (Zheng et al., 2010). Recently, two different groups generated and analyzed *iro3* loss-of-function mutants (Wang et al., 2020a; Wang et al., 2020b). Wang et al. (2020a) showed that the expression of Fe deficiency inducible genes increased in the root of *iro3* mutants, but Wang et al. (2020b) showed that OsIRO3 regulates only *OsNAS3*, but not other Fe deficiency inducible genes. Moreover, the underlying molecular mechanism by which OsIRO3 regulates Fe homeostasis remains unclear. In the present study, we showed that the loss-of-function of *OsIRO3* caused the up-regulation of many Fe deficiency inducible genes in the root. Further investigation found that OsIRO3 directly binds to and inhibits the promoter of *OsIRO2*. On the other hand, OsIRO3 physically interacts with OsPRI1/2 and represses the transactivation ability of the latter to OsIRO2. Additionally, OsIRO3 contains an EAR motif recruiting the OsTPL/OsTPRs corepressors, which partially accounts for its repression function.

## RESULTS

### Loss-of-function of *OsIRO3* impairs the Fe deficiency response

To further clarify the functions of OsIRO3 in the Fe deficiency response, we generated two *iro3* loss-of-function mutants with the CRISPR-Cas9 gene editing system. Two independent lines, *iro3-del1* with a deletion of nucleotide T in exon 4 and *iro3-del61* with a deletion of 61 bp in exon 3 were selected for further analysis (Figure 1A). Under Fe sufficient conditions, wild type and *iro3* mutant plants showed no discernable differences (Figure 1B). Under Fe deficient conditions, the wild type plants displayed the typical Fe deficiency symptom, chlorotic leaves. In contrast, three days after transfer to Fe deficiency medium, the mutants developed brown necrotic lesions in leaves, and the necrotic lesions gradually increased with the duration of Fe deficiency treatment. Meanwhile, compared with the wild type plants, the mutant plants developed dwarf shoots (Figure 1B). To explore whether loss-of-function of *OsIRO3* affects Fe homeostasis, we measured the Fe concentration in the root and shoot. Although the Fe concentration in the root and shoot of *iro3* mutants was not significantly different from that in the wild type under Fe sufficient conditions, the shoot Fe concentration in the *iro3* mutants was obviously higher than that in the wild type plants under Fe deficient conditions (Figure 1C), indicating that Fe translocation from root to shoot was enhanced in the *iro3* mutants. Collectively, these data suggest that loss-of-function of *OsIRO3* leads to the disruption of Fe homeostasis.

**Figure 1.**
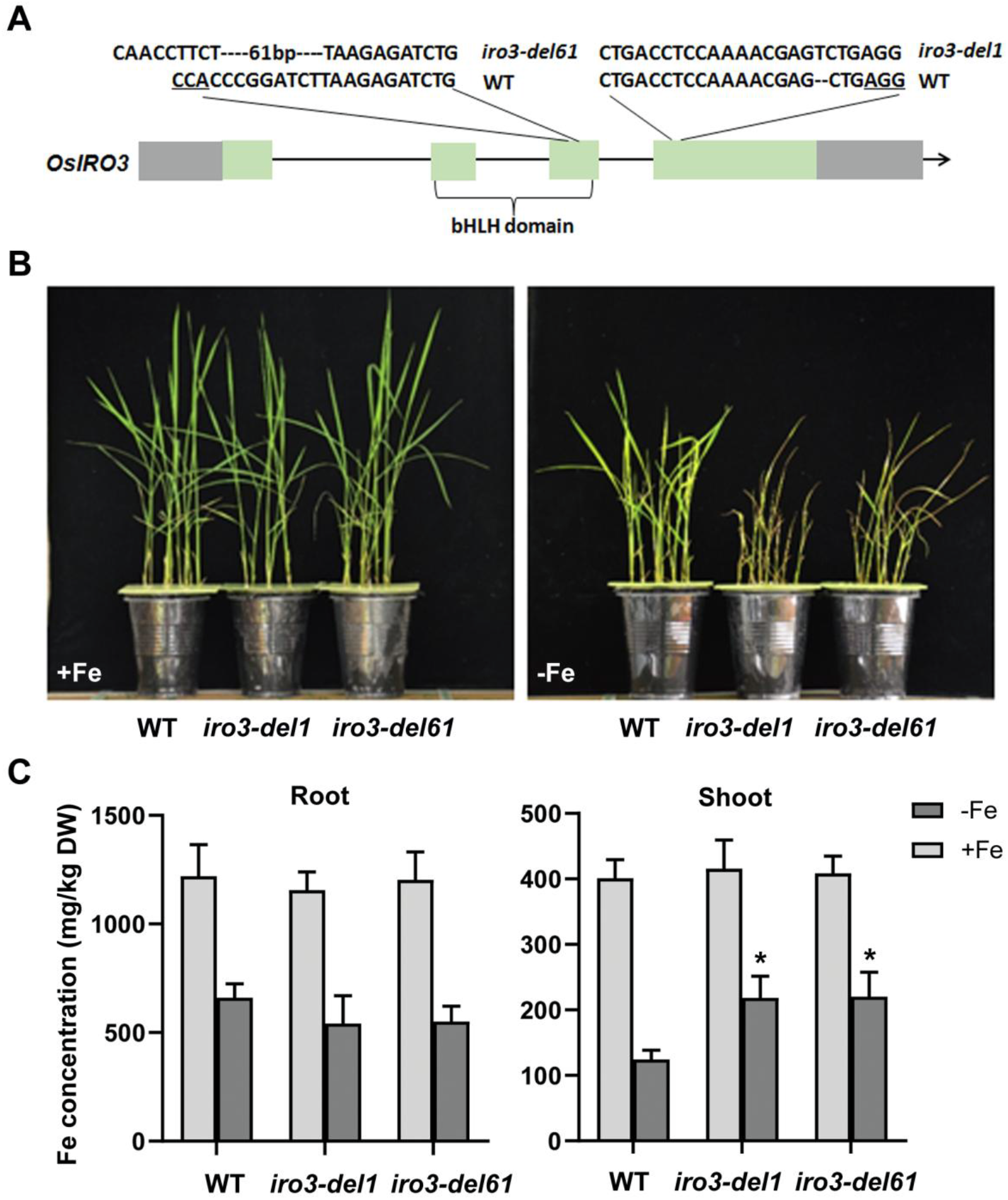
**Identification of *iro3* mutants.** (A) Mutations generated in the *iro3* mutants by CRISPR/Cas9. The underlined three letters indicate the PAM region. The *iro3-del1* mutant contains a deletion of nucleotide T in exon 4 and the *iro3-del61* mutant contains a deletion of 61 bp in exon 3. The genotypes of *iro3-del1* and *iro3-del61* are indicated. (B) Phenotypes of *iro3* mutants. Seeds were grown in +Fe (0.1 mM Fe^3+^) solution for two weeks, and then shifted to +Fe or –Fe (Fe free) solution for one week. (C) Fe concentration in the *iro3* mutants. Two-week-old seedlings grown in +Fe were transferred to +Fe or –Fe solution for 1 week. Shoots and roots were separately sampled and used for metal measurement. Error bars represent the SD (*n* = 3). The value which is significantly different from the corresponding wild-type (WT) value was indicated by * (*P* < 0.05), as determined by Student’s *t* test. DW, Dry weight.

### Loss-of-function of *OsIRO3* results in the activation of OsIRO2 regulon

Given that the loss of *OsIRO3* function disrupted the Fe homeostasis of rice, we wondered whether the expression of Fe deficiency inducible genes was changed in the *iro3* mutants. Therefore, we detected the gene expression of several representative Fe deficiency inducible genes. OsIRO2 and OsFIT are the master regulators of the Fe deficiency response, which positively regulate not only the Strategy II associated genes (Ogo et al., 2007; Liang et al., 2020; Wang et al., 2020), such as *OsNAS1*, *OsNAS2*, *OsNAAT1*, and *OsDMAS1* which encode the enzymes responsible for DMA synthesis (Inoue et al., 2003; Cheng et al., 2007; Bashir et al., 2017), *OsTOM1*, product of which accounts for the excretion of DMA (Nozoye et al., 2011), and *OsYSL15* which encodes an Fe (III)-DMA transporter (Inoue et al., 2009; Lee et al., 2009), but also the Strategy I associated gene *OsIRT1* (Ishimaru et al., 2006). When suffering Fe deficiency, rice plants initiate the expression of these genes. We found that the expression of *OsIRO2* and *OsFIT* and their downstream genes was considerably enhanced in the root of *iro3* mutants regardless of Fe status, which is in consistence with the negative function of OsIRO3. These results suggest that the expression of Fe deficiency inducible genes is disrupted in the *iro3* mutants.

### OsIRO3 directly represses the expression of *OsIRO2*

Given that *OsIRO2* and *OsFIT* and their downstream genes are downregulated in the *OsIRO3* overexpression plants (Zheng et al., 2010), and upregulated in the *iro3* plants (Figure 2), we speculated that OsIRO3 might directly regulate the expression of *OsIRO2* and *OsFIT.* The bHLH family transcription factors can bind to the E-box motifs of their target genes (Fisher and Goding, 1992). Several E-box motifs (CANNTG) exist in the promoters of *OsIRO2* and *OsFIT* (Figure S1; Zhang et al., 2017, 2020). Electrophoresis mobility shift assays (EMSAs) were performed to test whether OsIRO3 directly binds to the promoters of *OsIRO2* and *OsFIT*. 6xHis (histidine) tagged OsIRO3 (His-OsIRO3) was expressed and purified from *E. coli*. When His-OsIRO3 was incubated with biotin-labeled probe of *OsIRO2*, a prominent DNA-protein complex was detected. The binding capacity decreased as wild-type unlabeled probe increased, however, the addition of mutated wild-type unlabeled probe without an E-box did not affect the abundance of the DNA-protein complex (Figure 3A). The same EMSAs were conducted using the *OsFIT* probe, indicating that OsIRO3 could not bind to the promoter of *OsFIT.* These results suggest that OsIRO3 directly associates with the promoter of *OsIRO2*, but not of *OsFIT*.

**Figure 2.**
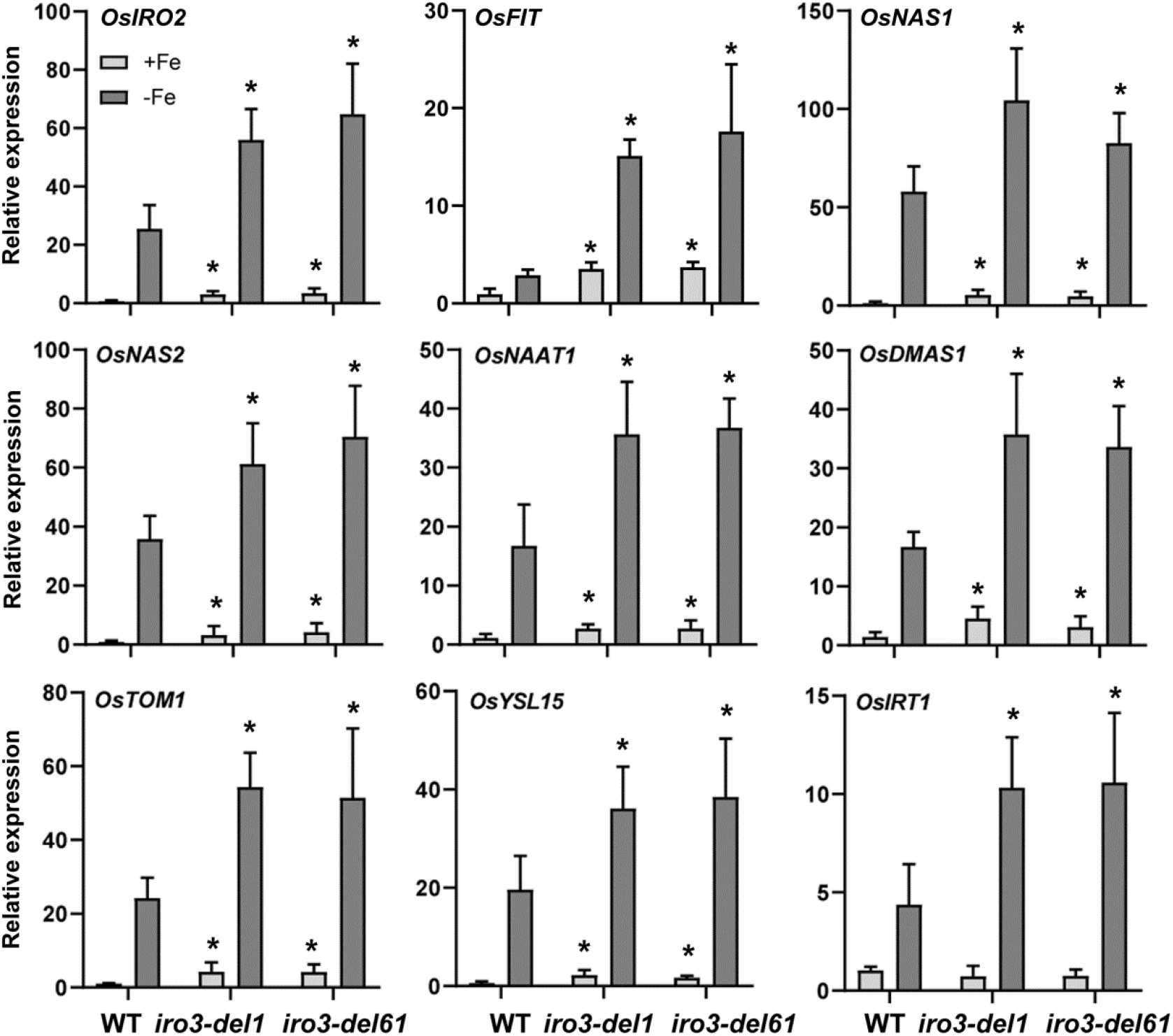
Expression of Fe deficiency inducible genes in the *iro3* mutants. Two-week-old seedlings grown in +Fe solution were transferred to +Fe or –Fe solution for 7 days. Roots were sampled and used for RNA extraction. The numbers above the bars indicate the corresponding mean values. Error bars represent the SD (*n* = 3). The value which is significantly different from the corresponding wild-type (WT) value was indicated by * (*P* < 0.05), as determined by Student’s *t* test.

**Figure 3.**
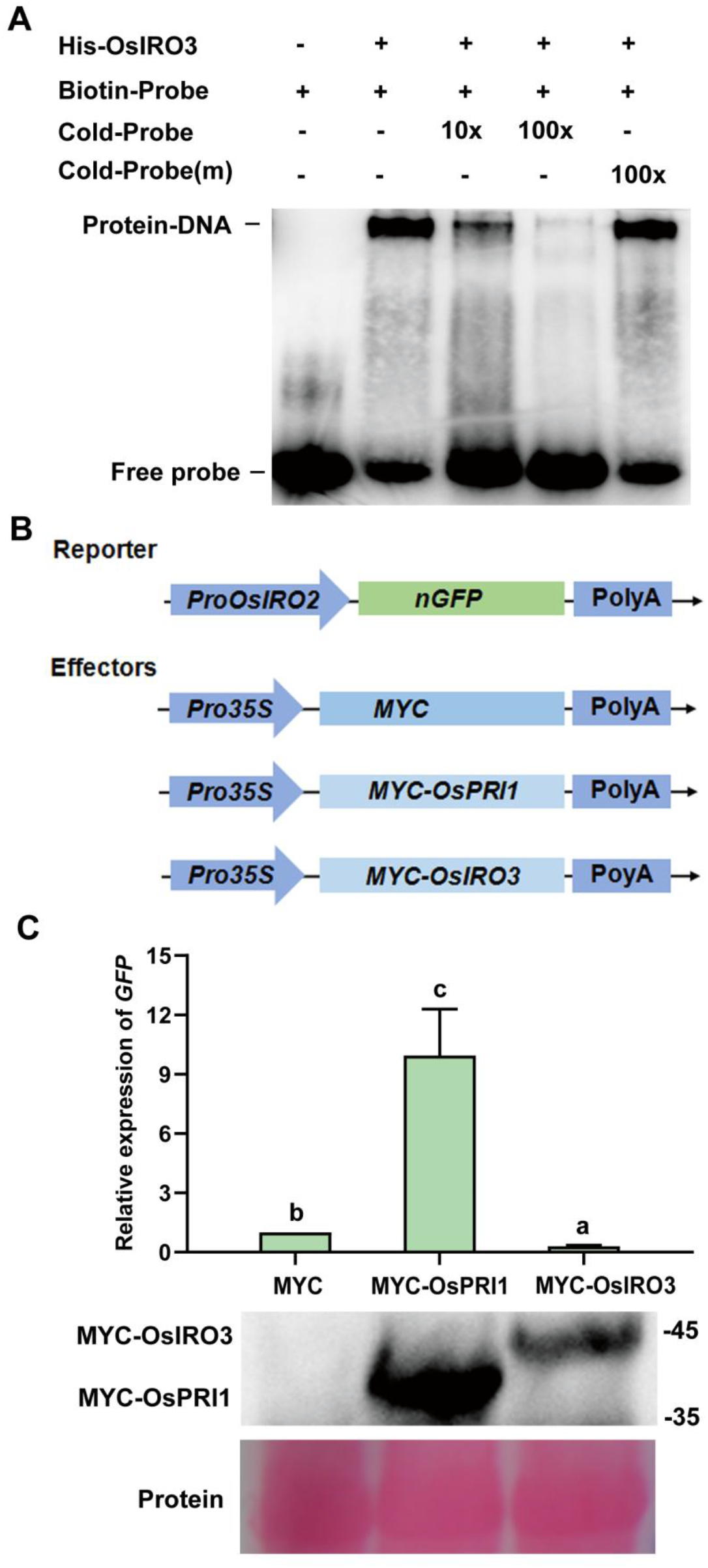
**OsIRO3 binds to the promoter of *OsIRO2*.** (A) EMSA assays. Biotin-labeled DNA probe was incubated with the recombinant His-OsIRO3 protein. An excess of unlabeled probe (Cold-Probe) or unlabeled mutated probe (Cold-Probe-m) was added to compete with labeled probe (Biotin-Probe). Biotin-probe incubated with His protein served as the negative control. (B) Schematic representation of the constructs used for transient expression assays. In the reporter, the *OsIRO2* promoter was used to drive a nuclear localization sequence fused GFP (nGFP). In the effectors, MYC, MYC-OsPRI1, and MYC-OsIRO3 are under the control of the cauliflower mosaic virus (CaMV) 35S promoter. (C) *GFP* transcript abundance. Protein levels of effectors were detected by immunoblot. Ponceau staining shows equal loading. *GFP* transcript abundance was normalized to *NPTII* transcript. The value with the empty vector (MYC) as an effector was set to 1. The different letters above each bar indicate statistically significant differences as determined by one-way ANOVA followed by Tukey’s multiple comparison test (*P* < 0.05).

To investigate whether OsIRO3 directly binds to and represses the promoter of *OsIRO2*, we prepared a reporter plasmid, *Pro_IRO2_:nGFP*, in which a nuclear localization signal fused *GFP* (*nGFP*) was driven by the 2204 bp upstream region of *OsIRO2* (Figure 3B). For the effector plasmids, MYC-tagged OsPRI1 and OsIRO3 were respectively cloned downstream of the 35S promoter. Transient expression assays were performed in tobacco leaves (Figure 3C). As a positive control, OsPRI1 significantly activated the expression of *GFP*. In contrast, OsIRO3 repressed the expression of *GFP*. These results indicate that OsIRO3 directly binds to and represses the *OsIRO2* promoter.

### OsIRO3 interacts with OsPRI1 and OsPRI2

Generally, bHLH transcription factors regulate downstream target genes by forming homodimers or heterodimers (Toledo-Ortiz et al., 2003). Considering that the Fe deficiency inducible genes regulated by OsIRO3 are also regulated by OsPRIs (OsPRI1, OsPRI2, and OsPRI3) (Zhang et al., 2017; Zhang et al., 2020), we speculated that OsIRO3 interacts with OsPRIs to form heterodimers to modulate the Fe deficiency response.

Yeast two-hybrid assays were used to test the potential protein interactions. Since the strong self-activation of the full length OsIRO3, the N-terminal part of OsIRO3 (OsIRO3n) containing the bHLH domain was fused with the GAL4 DNA binding domain (BD) as the bait. Four OsPRIs were respectively fused to the GAL4 activating domain (AD) as preys. Yeast two-hybrid assays showed that OsPRI1 and OsPRI2, but not OsPRI3 and OsPRI4, interact with OsIRO3 (Figure 4A). To further verify the interactions between OsIRO3 and OsPRI1/2, pull-down assays were carried out. OsPRI1 and OsPRI2 were fused with the GST (glutathione S-transferase) tag respectively, and OsIRO3 was fused with the 6xHis tag. Proteins were expressed and purified from *E. coli*. GST, GST-OsPRI1, and GST-OsPRI2 were respectively co-incubated with His-tagged OsIRO3 and then eluted. The immunoblot results showed that GST-OsPRI1 and GST-OsPRI2 pulled down His-OsIRO3, but GST did not (Figure 4B). To further verify whether their interactions also occur in plant cells, tripartite split-GFP assays were performed in *Nicotiana benthamiana* leaves. The GFP10 fragment was fused with the N-end of OsIRO3 (GFP10-OsIRO3) and the GFP11 fragment with the C-end of OsPRI1/2 (OsPRI1/2-GFP11). W hen OsPRI1-GFP11 (or OsPRI2-GFP11) was co-expressed with GFP10-OsIRO3 and GFP1-9, strong fluorescence signal was detected in the nucleus, whereas fluorescence signal was hardly detected in the cells co-expressing GFP11, GFP10-OsIRO3, and GFP1-9 (Figure 4C). All these results indicate that OsIRO3 interacts with OsPRI1 and OsPRI2.

**Figure 4.**
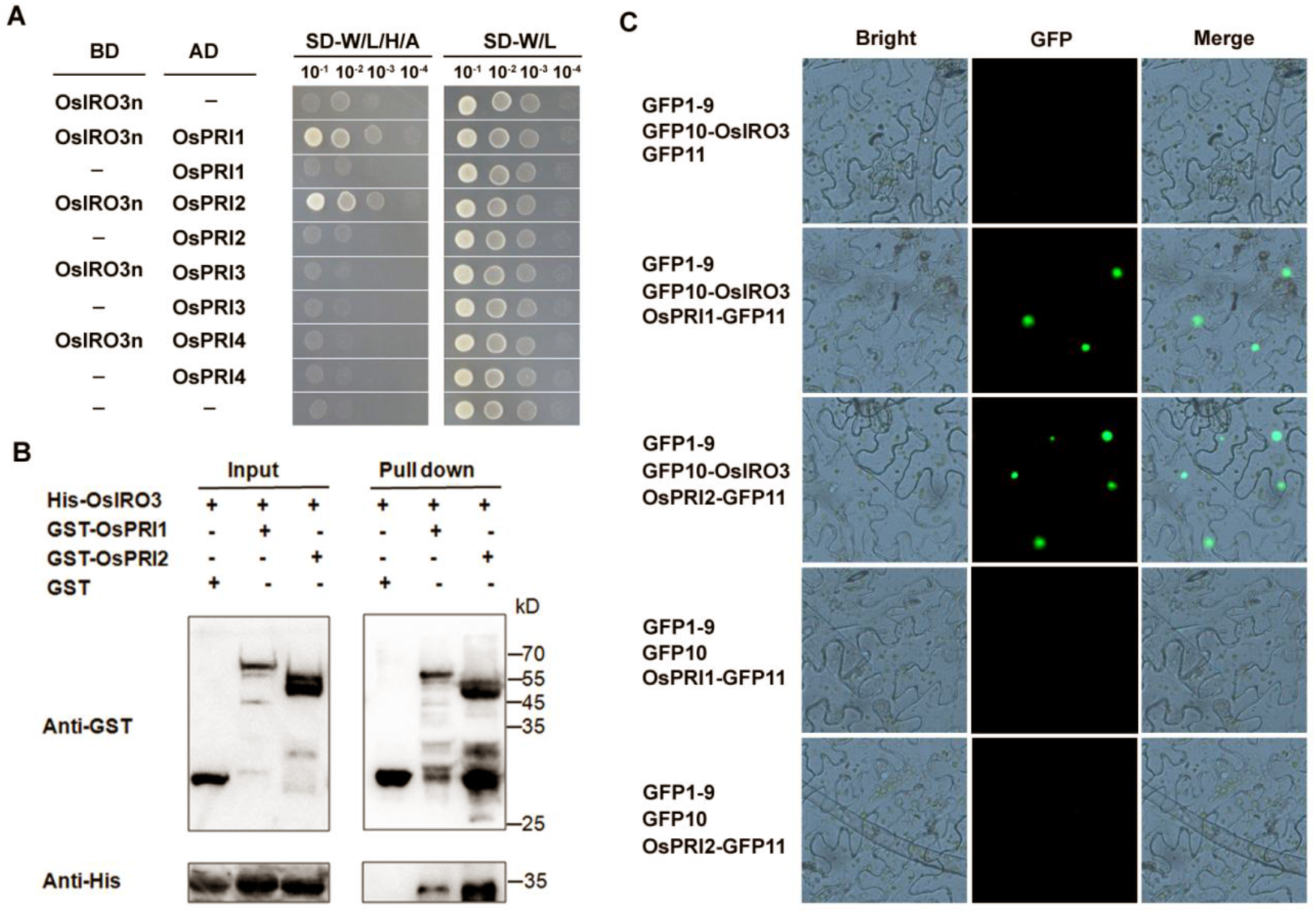
**OsIRO3 physically interacts with OsPRI1 and OsPRI2.** (A) Yeast two-hybrid analysis of the interactions between OsIRO3 and OsPRI1/2. Yeast cotransformed with different BD and AD plasmid combinations was spotted on synthetic dropout medium lacking Leu/Trp (SD-W/L) or Trp/Leu/His/Ade (SD-W /L/H/A). (B) Pull-down assays. OsPRI1/2 were respectively fused with the GST tag, and OsIRO3 was fused with the His tag. Recombinant proteins were expressed in *E. coli*. Proteins were pulled down by glutathione Sepharose 4B and detected using the anti-His or anti-GST antibody. (C) Protein interactions of OsIRO3 and OsPRI1/2 in plant cells. Tripartite split-sfGFP complementation assays were performed. OsPRI1 and OsPRI2 were respectively fused with GFP11, and OsIRO3 with GFP10. The constructs were introduced into agrobacterium respectively, and the indicated combinations were co-expressed in *N. benthamiana* leaves.

### OsIRO3 inhibits the transactivation of OsPRI1 towards *OsIRO2*

It has been established that OsPRI1/2/3 positively regulate the expression of *OsIRO2* through directly binding to and activating its promoter (Zhang et al., 2017, 2020). Considering that OsIRO3 interacts with OsPRI1/2, we wanted to know whether OsIRO3 interferes with the transactivation ability of OsPRI1/2 towards *OsIRO*2. We carried out transient expression assays using the reporter-effector system above-mentioned (Figure 3B). Compared with the control effector (MYC), co-expression of OsIRO3 with OsPRI1 significantly weakened the GFP signal (Figure 5A). These data suggest that OsIRO3 inhibits the transactivation of OsPRI1 towards *OsIRO2*.

**Figure 5.**
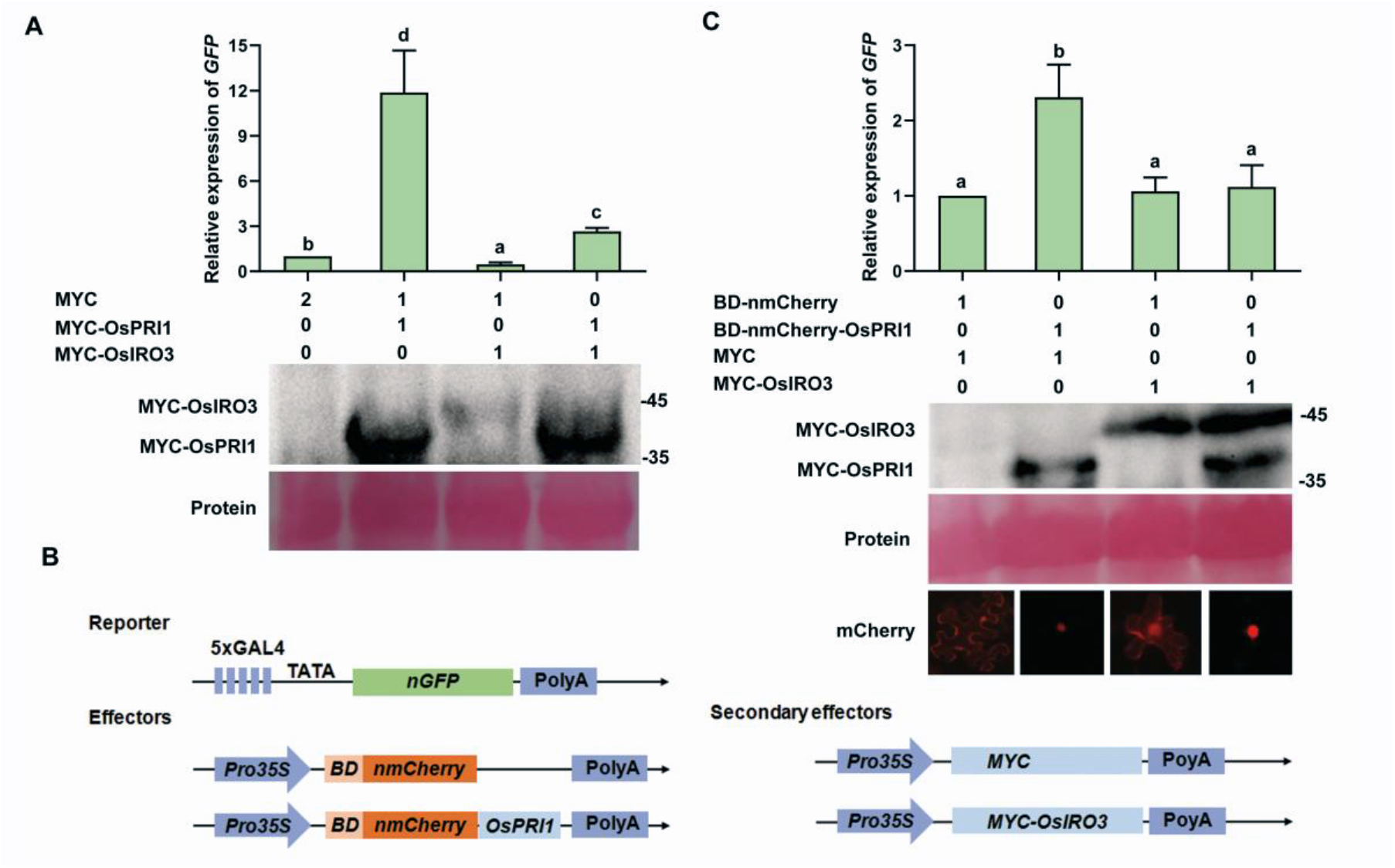
**OsIRO3 antagonizes the transcriptional activation ability of OsPRI1.** (A) OsIRO3 represses the transcription activation of OsPRI1. The reporter and effectors are shown in Figure 3B. Protein levels of effectors were detected by immunoblot. Ponceau staining shows equal loading. The *GFP/NPTII* ratio represents the *GFP* levels relative to the internal control *NPTII*. (B) Schematic representation of the constructs used for transient expression assays. In the reporter, five repeats of GAL4 binding motif and the minimal CaMV 35S promoter was used as the promoter to drive the nGFP. In the effectors, BD-nmCherry and BD-nmCherry-OsPRI1 are under the control of 35S promoter. In the secondary effectors, MYC and MYC-OsIRO3 are under the control of 35S promoter. (C) OsIRO3 inhibits the transcriptional activation ability of OsPRI1 by direct protein-protein interaction. Protein levels of effectors were detected by immunoblot. Ponceau staining shows equal loading. The abundance of *GFP* was normalized to that of *NPTII*. The value with the control (nmCherry) was set to 1. The different letters above each bar indicate statistically significant differences as determined by one-way ANOVA followed by Tukey’s multiple comparison test (*P* < 0.05).

It has been confirmed that OsPRI1 and OsPRI2 also bind to the *OsIRO2* promoter, raising the possibility that OsIRO3 competes with OsPRI1/2 for binding to the *OsIRO2* promoter, hence reducing the expression of *OsIRO2*. To further clarify whether OsIRO3 directly represses the transactivation function of OsPRI1 by protein interaction, we employed the GAL4-based reporter-effector system (Li et al., 2022). For the reporter, the *nGFP* was driven by a synthetic promoter which consists of five repeats of GAL4 binding motif and the minimal CaMV 35S promoter (Figure 5B). For the effector, the GAL4 BD fused with an NLS-mCherry and OsPRI1 was driven by the 35S promoter (Figure 5B). Compared with the control (nmCherry), OsPRI1 activated the expression of *GFP* (Figure 5C). W hen OsIRO3 was co-expressed with OsPRI1, the expression of *GFP* was significantly suppressed. These data suggest that OsIRO3 can inhibit the transactivation of OsPRI1 through the direct protein interaction with OsPRIs.

### OsIRO3 interacts with the co-repressors OsTPL/OsTPRs

Many negative transcription factors exert repression functions by their EAR recruiting transcriptional co-repressors TOPLESS/TOPLESS-RELATED (TPL/TPRs). Two types of EAR motifs, LxLxL and DLNxxP, have been characterized (Kagale et al., 2010; Causier et al., 2012). We searched all known transcription factors involved in the Fe deficiency response of rice for EAR motifs, finding that both OsIRO2 and OsIRO3 contain a typical LxLxL EAR motif (Figure 6A).

**Figure 6.**
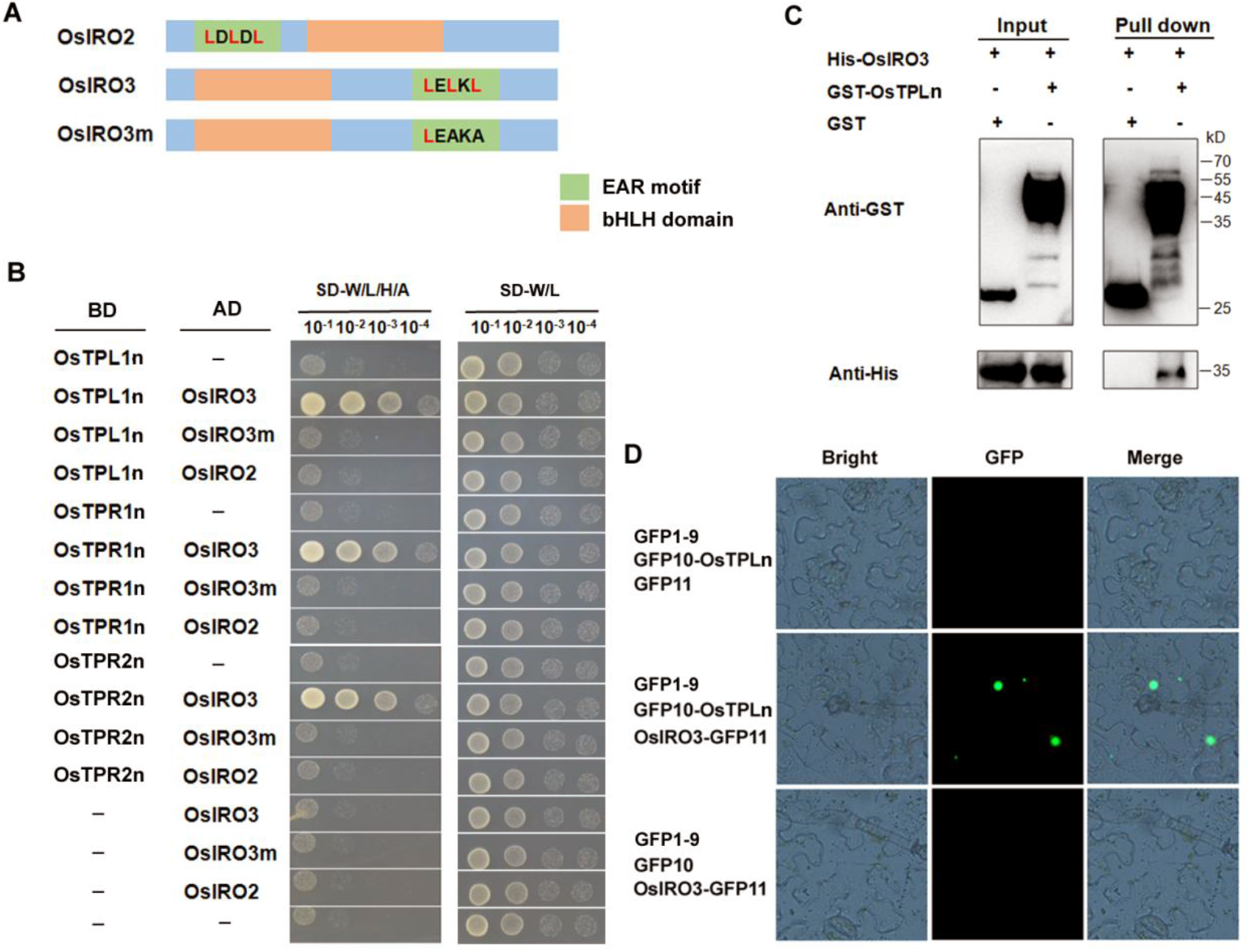
**OsIRO3 interacts with the co-repressors OsTPL/OsTPRs.** (A) Schematic diagram of bHLH domain and EAR motif in the OsIRO2 and OsIRO3. (B) The EAR motif is required for the interactions between OsIRO3 and OsTPL/OsTPRs. Yeast cotransformed with different BD and AD plasmid combinations was spotted on synthetic dropout medium lacking Leu/Trp (SD-T/L) or Trp/Leu/His/Ade (SD-T/L/H/A). (C) Pull-down assays. The N-terminal of OsTPL was fused with the GST tag, and OsIRO3 was fused with the His tag. Recombinant proteins were expressed in *E. coli*. Proteins were pulled down by glutathione Sepharose 4B and detected using the anti-His or anti-GST antibody. (D) Interaction of OsIRO3 and OsTPLn in plant cells. Tripartite split-sfGFP complementation assays were performed. OsTPLn was fused with GFP10, and OsIRO3 with GFP11. The constructs were introduced into agrobacterium respectively, and the indicated combinations were co-expressed in *N. benthamiana* leaves.

Subsequently, we wondered whether both OsIRO2 and OsIRO3 could interact with OsTPL/OsTPRs. We employed the yeast two-hybrid assays to test their protein interactions (Figure 6B). Given that the N-terminal of OsTPL/OsTPRs is responsible for the interaction with EAR motifs, three N-terminal truncated OsTPLn, OsTPR1n, and OsTPR2n were respectively fused with the BD. The full-length of OsIRO2 and OsIRO3 were respectively fused with the AD. The results showed that OsIRO3, but not OsIRO2, could interact with OsTPL/OsTPRs, which is consistent with the fact that OsIRO3 is a negative regulator and OsIRO2 a positive regulator. To further investigate whether the EAR motif of OsIRO3 is responsible for the interactions with OsTPL/OsTPRs, we constructed a mutated version of OsIRO3 (OsIRO3m) with a mutated EAR motif (LxAxL). Interaction tests indicated that the mutation of EAR enabled OsIRO3m not to interact with OsTPL/OsTPRs, suggesting that the EAR motif is required for the interactions. Next, we performed pull-down assays in which OsTPLn was used as a representative (Figure 6C). The results suggest that OsTPLn could pull down OsIRO3. The tripartite split-GFP assays further confirmed that their interaction occurs in the nucleus (Figure 6D). Taken together, these results indicated that OsIRO3 interacts with OsTPL/OsTPRs co-repressors and its EAR motif is responsible for the interactions.

### The repression function of OsIRO3 partially depends on its EAR motif

Having confirmed that OsIRO3 interacts with OsTPL/OsTPRs through its EAR motif, we asked if the EAR motif is crucial for the repression function of OsIRO3. To test this, we carried out reporter-effector transient expression assays, in which *Pro_OsIRO2_:nGFP* was used as the reporter. OsIRO3 strongly reduced the expression of *GFP* whereas OsIRO3m displayed weak inhibitory effect on the expression of *GFP*. We further examined the influence of OsIRO3m on OsPRI1. When co-expressed with OsPRI1, both OsIRO3 and OsIRO3m repressed the expression of *GFP* compared with the control (MYC), but the inhibitory effect of OsIRO3m was not as strong as that of OsIRO3 (Figure 7A). These data suggest that the repression function of OsIRO3 is partially dependent on its EAR motif.

**Figure 7.**
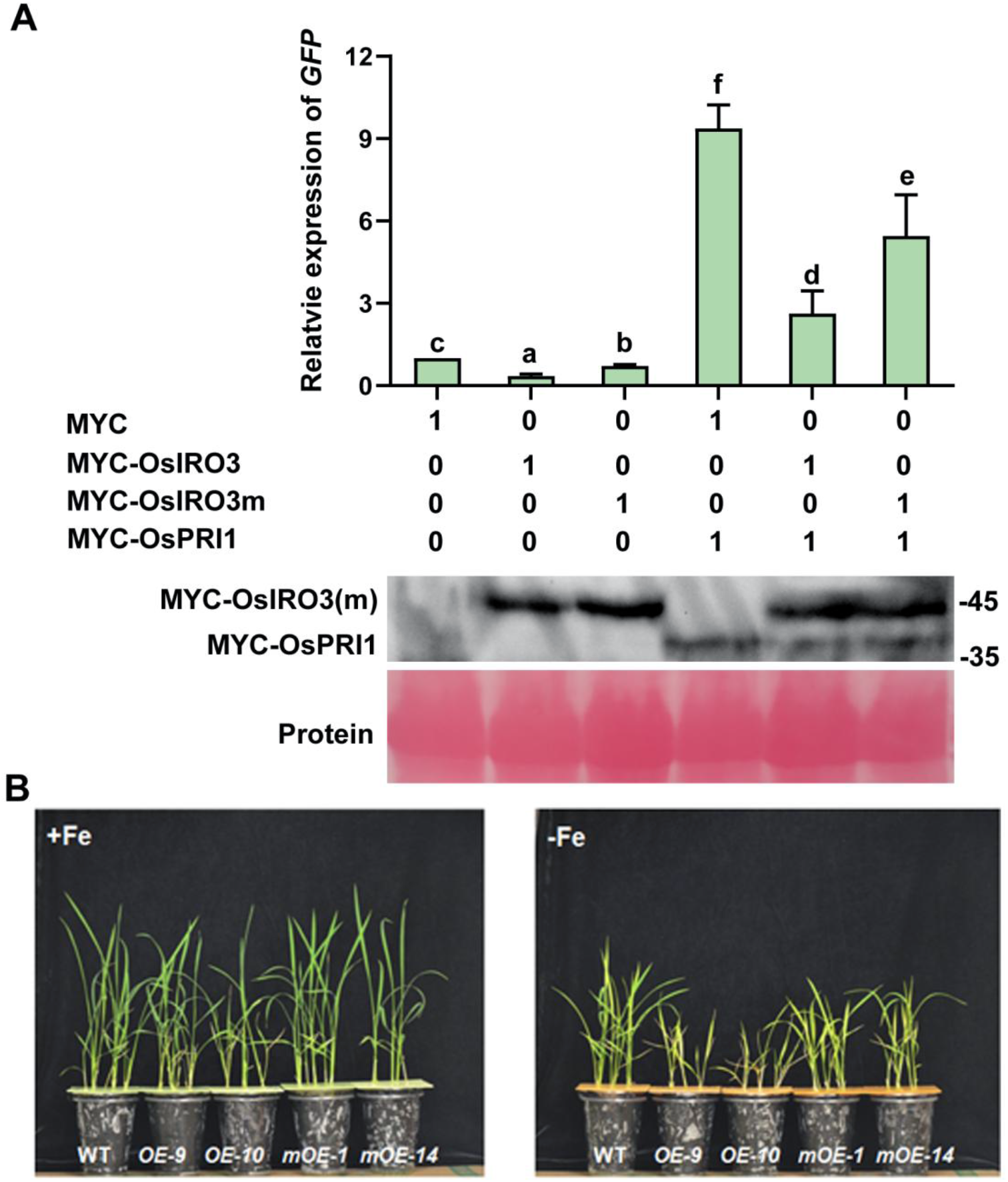
**The EAR motif partially contributes to the repression function of OsIRO3.** (A) The EAR motif is partially required for the repression function of OsIRO3. The reporter and effectors are shown in Figure 3B. Protein levels of effectors were detected by immunoblot. Ponceau staining shows equal loading. The abundance of *GFP* was normalized to that of *NPTII*. The value with the empty vector as an effector was set to 1. The different letters above each bar indicate statistically significant differences as determined by one-way ANOVA followed by Tukey’s multiple comparison test (*P* < 0.05). (B) Phenotypes of *OsIRO3*(*m*) overexpression plants. Seeds were germinated on wet paper for seven days, and then seedlings were shifted in +Fe (0.1 mM Fe^3+^) or –Fe (Fe free) solution for two weeks.

Given that the EAR motif affects the repression function of OsIRO3, we wanted to know whether the EAR motif also affects its biological functions. For this aim, we constructed transgenic plants overexpressing *OsIRO3* and *OsIRO3m,* respectively (Figure S2). Under Fe sufficient conditions, both *OsIRO3-OX* and *OsIRO3m-OX* plants grew as well as the wild type plants (Figure 7B). Under Fe deficient conditions, two independent *OsIRO3* overexpression lines (*OE9* and *OE10*) showed hypersensitivity to Fe deficiency compared with the wild-type plants, including chlorotic leaves and reduced shoot height, which is consistent with the previous study (Zheng et al., 2010). Although the *OsIRO3m* overexpression lines (*mOE-1* and *mOE-14*) also displayed sensitivity to Fe deficiency, they were less sensitive to Fe deficiency compared with the *OsIRO3* overexpression plants. Taken together, our results suggest that the EAR motif is necessary for the biological functions of OsIRO3.

## DISCUSSION

Plants have evolved intricate mechanisms to maintain Fe homeostasis. When facing Fe deficiency conditions, plants up-regulate the expression of Fe deficiency inducible responsive genes, thereby promoting Fe absorption to meet the plant’s needs. However, excessive Fe uptake is prone to result in reactive oxygen species which are toxic to plant cells. The balance between positive regulatory factors to activate the Fe uptake system and negative regulatory factors to suppress it maintains Fe homeostasis in plants. OsIRO2 is a crucial regulator for the Fe uptake system in rice. To maintain Fe homeostasis, rice plants activate *OsIRO2* under Fe deficient conditions, and suppress it under Fe sufficient conditions. Previous studies have shown that three OsPRI proteins directly and positively regulate *OsIRO2* (Zhang et al., 2017, 2020). However, it is still unclear which transcription factors directly and negatively regulate *OsIRO2*. Here, we provide evidence that OsIRO3 not only directly represses the expression of *OsIRO2* by associating with its promoter, but also indirectly by inhibiting the transcription activation of OsPRI1/2 to *OsIRO2*.

OsIRO3 was identified as a negative regulator of Fe homeostasis since its overexpression leads to chlorotic leaves, decreased Fe concentration and reduced expression of many Fe deficiency inducible genes (Zheng et al., 2010). Notably, the loss-of-function of *OsIRO3* causes enhanced shoot Fe accumulation and necrotic spots in leaves under Fe deficient conditions (Figure 1B). Two different group recently reported the similar phenotypes of *iro3* mutants and explained that the increased ROS might contribute to the leaf necrosis of *iro3* mutants (Wang et al., 2020a; Wang et al., 2020b).

However, regarding to the expression of Fe deficiency inducible genes, these two groups showed different results. Wang et al. (2020a) showed that the expression of Fe deficiency inducible genes was increased in *iro3* under Fe deficient conditions, and Wang et al. (2020b) did not observe the change of those genes except for *OsNAS3*. It is very likely that the different results are attributed to their different experimental conditions. Our results support that loss-of-function of *OsIRO3* promotes the expression of Fe deficiency inducible genes.

When suffering Fe deficiency, plants stimulate their Fe uptake systems to acquire more Fe. At the same time, Fe deficiency can lead to the inactivity of many ROS scavengers which require Fe as co-factors. A burst of Fe influx is prone to production of radical oxygen species, which are toxic to plant cells. The finetune of Fe uptake system ensures the viability of cells, hence the health of plants. As a key regulator of the Fe uptake system, the transcription of *OsIRO2* must be tightly regulated. It has been revealed how the transcription of *OsIRO2* is activated directly under Fe deficient conditions (Zhang et al., 2017, 2020). It was unclear how the transcription of *OsIRO2* is repressed directly. Our data suggest that OsIRO3 directly recognizes the *OsIRO2* promoter (Figure 3A), and represses the transcription of *OsIRO2* (Figure 3C). In addition to the direct repression, OsIRO3 also indirectly represses the transcription of *OsIRO2* since OsIRO3 interacts with OsPRI1/2 to attenuate their transactivation activity towards *OsIRO2* (Figure 5A). Thus, the balance between promotion and repression of *OsIRO2* finetunes the abundance of *OsIRO2*, hence maintaining appropriate Fe levels in cells. In the *iro3* mutants, the repression of *OsIRO2* by OsIRO3 is cancelled, and the expression of *OsIRO2* is out of control, finally resulting in the Fe toxicity symptom of leaves. It has been confirmed that OsIRO2 and OsFIT interact with each other to control the Strategy II genes (Liang et al., 2020; Wang et al., 2020). Notably, the overexpression of *OsFIT* also causes leaf necrosis symptoms similar to that of *iro3* under Fe deficient conditions (Liang et al., 2020). Therefore, it is very likely that the excessive activation of Fe uptake system accounts for the Fe toxicity symptoms under Fe deficient conditions. Meanwhile, the *iro3* mutants accumulate more Fe only in the shoot under Fe deficient conditions (Figure 1C), and *OsIRO2* and *OsFIT* and their target genes are induced in the roots of *iro3* under Fe deficient conditions (Figure 2). It is very likely that OsIRO2 controls not only the Fe uptake from soil to root, but also Fe translocation from root to shoot. Although *OsFIT* is also negatively regulated by OsIRO3, a direct link between them is still missing. OsIRO3 more likely indirectly down-regulates its transcription.

OsIRO3 is a negative regulator of Fe homeostasis, however, it was unclear how OsIRO3 exerts its repressive function. There are two types of transcriptional repressors, active and passive repressors. Active transcriptional repressors function by recruiting the transcriptional corepressors, such as OsTPL/OsTPRs, while passive repressors compete with positive transcription factors for binding to target gene promoters. Here, we reveal that OsIRO3 can act as an active repressor by recruiting the transcriptional corepressors OsTPL/OsTPRs (Figure 6). Our EMSA assays confirmed that OsIRO3 can bind to the *OsIRO2* promoter which is also targeted by OsPRI1/2/3 (Zhang et al., 2017, 2020). It is very likely that OsIRO3 competes with OsPRI1/2/3 for binding to the *OsIRO2* promoter so that less OsPRI proteins are involved in the transcription initiation of *OsIRO2*. Therefore, OsIRO3 might also act as a passive repressor.

OsIRO2 is an ortholog of Arabidopsis bHLH Ib subgroup members (AtbHLH38, AtbHLH39, AtbHLH100, and AtbHLH101). Arabidopsis bHLH Ib subgroup members interact with AtFIT to modulate the expression of Strategy I genes (Yuan et al., 2008; Wang et al., 2013) while OsIRO2 interacts with OsFIT to control the expression of Strategy II genes (Liang et al., 2020; Wang et al., 2020). Arabidopsis bHLH IVc proteins directly regulate bHLH Ib genes (Zhang et al., 2015; Li et al., 2016; Liang et al., 2017) while rice bHLH IVc proteins directly target *OsIRO2* (Zhang et al., 2017, 2020). Although plants utilize different strategies to take up Fe from soil, they have evolved this conserved regulatory mechanism to control the Fe uptake systems. We reveal that OsIRO3 functions as a brake to restrict the expression of *OsIRO2* under Fe deficient conditions. A latest study revealed that AtbHLH11 interacts with bHLH IVc members and inhibits the transcription activation of the latter to bHLH Ib genes, and that AtbHLH11 can also recruit the co-repressors AtTPL/AtTPRs (Li et al., 2022). However, there is no evidence supporting that bHLH Ib genes are the direct targets of AtbHLH11. Furthermore, *OsIRO3* is induced under Fe deficient conditions (Zheng et al., 2010) whereas *AtbHLH11* is repressed (Li et al., 2022). Thus, OsIRO3 regulates the Fe deficiency response in a manner different from AtbHLH11. OsIRO3 is a close homolog of Arabidopsis AtPYE (POPEYE/AtbHLH47) (Zheng et al., 2010). AtPYE directly targets *AtZIF1, AtFRO3* and *AtNAS4* which are involved in Fe homeostasis (Long et al., 2010). Similarly, OsIRO3 directly regulates *OsNAS3* in rice (Wang et al., 2020b). Arabidopsis bHLH IVc subgroup members (AtbHLH34, AtbHLH104, AtbHLH105, and AtbHLH115) correspond to rice bHLH IVc subgroup members (OsPRI1, OsPRI2, OsPRI3 and OsPRI4) (Zhang et al., 2020). Similar to AtPYE which physically interacts with three bHLH IVc members AtbHLH104/105/115 (Long et al., 2010; Selote et al., 2015), OsIRO3 interacts with two bHLH IVc members OsPRI1/2 (Figure 4). Although AtPYE and OsIRO3 share these similarities, they regulate Fe homeostasis in different manners. Unlike the *iro3* mutant plants which accumulate more Fe only in the shoot under Fe deficient conditions, *pye* mutant plants accumulate more Fe both in the root and shoot irrespective of Fe status (Long et al., 2010). Loss-of-function of *AtPYE* does not affect the expression of Fe uptake genes, such as *AtIRT1* and *AtFRO2* (Long et al., 2010), whereas loss-of-function of *OsIRO3* facilitates the expression of Fe uptake genes. Thus, there is a functional divergence between OsIRO3 and AtPYE, and Arabidopsis and rice have developed different regulatory mechanisms to repress bHLH Ib genes.

This study expands our knowledge of the Fe homeostasis transcription network mediated by the OsIRO3-OsIRO2 module. Based on our findings, we propose a putative working model for OsIRO3 (Figure 8). The transcription of *OsIRO2* is regulated positively by OsPRIs (OsPRI1 and OsPRI2), but negatively by OsIRO3. OsPRIs directly associate with and activate the promoter of *OsIRO2*. In contrast, OsIRO3 represses *OsIRO2* in two different manners: (a) directly binding to the *OsIRO2* promoter and repressing its transcription by recruiting the co-repressors OsTPL/OsTPRs; (b) inhibiting the transcription activation ability of OsPRIs towards *OsIRO2*. Under Fe deficient conditions, the balance of OsPRIs and OsIRO3 ensures that *OsIRO2* is expressed at an appropriate level. When the function of *OsIRO3* is lost, the repression of *OsIRO2* is removed, resulting in excessive expression of *OsIRO2*. This work enhances our understanding of the Fe deficiency response signaling pathway in rice.

**Figure 8.**
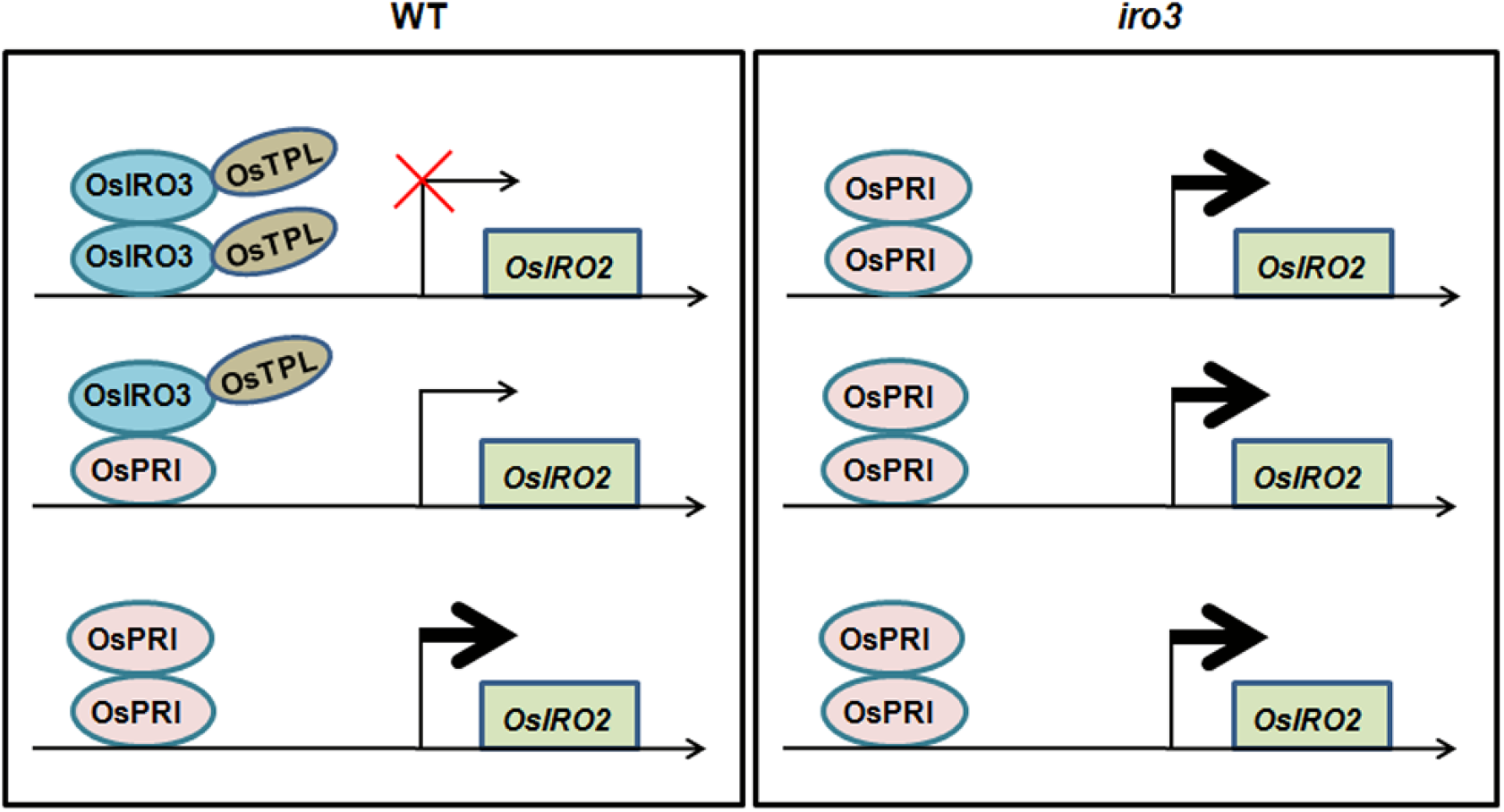
A working model of OsIRO3 in Fe homeostasis. Like OsPRIs (OsPRI1/2), OsIRO3 directly associates with the promoter of *OsIRO2*. OsIRO3 can function as an active repressor by recruiting the transcriptional corepressors OsTPL/OsTPRs. OsIRO3 also physically interacts with OsPRIs. Under Fe-deficient conditions, OsIRO3 and OsPRIs are abundant and antagonistically regulate the expression of *OsIRO2*. In wild type (WT), OsPRIs activate the expression of *OsIRO2*; on the other hand, OsIRO3 represses the expression of *OsIRO2* either by directly binding to the promoter of *OsIRO2* or by inhibiting the transcription activation of OsPRIs towards *OsIRO2*. The balance of OsPRIs and OsIRO3 under Fe deficient conditions ensures an appropriate level of *OsIRO2*. In the *iro3* mutants, the repression of *OsIRO2* by OsIRO3 is cancelled, resulting in the over-accumulation of *OsIRO2* transcripts.

## MATERIALS AND METHODS

### Plant materials and growth conditions

Rice cultivar ‘Nipponbare’ was used in this study. Plants were grown in a greenhouse with a photoperiod of 14 h light 28°C and 10 h dark at 22°C. For hydroponic culture assays, Fe-sufficient solution was prepared in half-strength Murashige and Skoog with 0.1 mM Fe (III)-EDTA and Fe-deficient solution in the same media without Fe.

### Generation of transgenic plants

The editing vectors were constructed as described previously (Liang et al., 2016). Two different target sites for *OsIRO3* were designed. The OsU6a promoter driving the sgRNA containing a specific target site was cloned into the pMH-SA vector by the restriction enzyme sites *Spe* I and *Asc* I. Two independent constructs were used for rice transformation. Homozygous mutant lines were identified by PCR sequencing.

For the construction of overexpression vectors, HA-OsIRO3 and HA-OsIRO3m were amplified from GAD-OsIRO3 and GAD-OsIRO3m, respectively, and cloned between the maize ubiquitin promoter and the NOS terminator in the pUN1301 binary vector.

### EMSA

OsIRO3 was cloned into the pET-28a(+) vector and the resulting plasmids was introduced into *Escherichia coli* BL21(DE3) for protein expression. Cultures were incubated with 0.5 M isopropyl β-D-1-thiogalactopyranoside at 22°C for 16h, and proteins were extracted and purified by using the His-tag Protein Purification Kit (Beyotime, China) following the manufacturer’s protocol. EMSA was performed using the Chemiluminescent EMSA Kit (Beyotime, China) following the manufacturer’s protocol. Briefly, two complementary single-strand DNA primers were was synthesized with a biotin label at the 5’ end. Two complementary primers were mixed and annealed to form the biotin-probe. The two biotin-unlabeled single-strand DNA primers were used as competitors, and the His protein alone was used as the negative control.

### Reverse transcription and quantitative PCR

Total RNA extracted from rice roots using TRIzol reagent (Invitrogen, USA) . cDNA was synthesized by the use of PrimeScript™ RT reagent Kit with gDNA Eraser (Perfect Real Time) according to the reverse transcription protocol (Takara). The resulting cDNA was subjected to relative quantitative PCR using a SYBR Premix Ex Taq^TM^ kit (TaKaRa) on a Roche LightCycler 480 real-time PCR machine, according to the manufacturer’s instructions. All PCR amplifications were performed in three biological replicates with *OsACTIN1* and *OsOBP* as the internal controls. Primers used in this paper are listed in Supplemental Table S1.

### Fe Measurement

To determine Fe concentration, 14-d-old seedlings grown in 1/2 MS liquid with 0.1 mM Fe (III)-EDTA were transferred to Fe-sufficient (0.1 mM Fe (III)-EDTA) or Fe-deficient (Fe free) liquid media for 7 d. The shoots and roots were harvested separately and dried at 65°C for 3 d. For each sample, about 500 mg dry weight of roots or shoots was digested with 5 mL of 11 M HNO_3_ and 2 mL of 12 M H_2_O_2_ for 30 min at 220°C. Fe concentration was measured using Inductively Coupled Plasma Mass Spectrometry (ICP-MS).

### Yeast two-hybrid assays

For yeast two-hybrid assays, the N-terminal truncated version of OsIRO3n, OsTPLn, OsTPR1n, and OsTPR2n were respectively cloned into pGBKT7. The sequence encoding full-length OsPRI1, OsPRI2, OsIRO2, OsIRO3, and OsIRO3m were respectively cloned into pGADT7. Vectors were transformed into yeast strain Y2HGold (Clontech, Japan). Growth was determined as described in the Yeast Two-Hybrid System User Manual (Clontech, Japan).

### Protein interaction in plant cells

The GFP1-9, GFP10, and GFP11 sequences of superfolder GFP were cloned into separate pER8 vectors under the estradiol induction promoter, generating pTG-GFP1-9, pTG-GFP10, and pTG-GFP11, respectively. OsIRO3 and OsTPLn were cloned into pTG-GFP10 with an N-terminal GFP10 tag, and OsPRI1/2 and OsIRO3 were cloned into pTG-GFP11 with a C-terminal GFP11 tag. All vectors were introduced into *A. tumefaciens* (strain EHA105) and various combinations of *Agrobacterium* cells were infiltrated into leaves of *N. benthamiana* in infiltration buffer (0.2 mM acetosyringone, 10 mM MgCl_2_, and 10 mM MES [pH 5.6]). Gene expression was induced 1 day after agroinfiltration by injecting 20 mM β-estradiol into the abaxial side of the leaves. Epidermal cells were observed and recorded under a Carl Zeiss Microscope.

### Pull-down assays

OsPRI1/2 and OsTPLn were cloned into pGEX-4T-1 respectively, and OsIRO3 was cloned into pET-28a (+). All plasmids were introduced into *Escherichia coli* BL21 cells (TransGen Biotech). GST, GST-OsPRI1/2, GST-OsTPLn, and His-OsIRO3 proteins were induced by 0.1 mM isopropyl-b-thiogalactopyranoside (IPTG) at 16°C for 20 h. Soluble GST, GST-OsPRI1/2, and GST-OsTPL-N were extracted and immobilized to glutathione affinity resin (Beyotime Biotechnology). For pull-down assays, His-OsIRO3 fusion proteins purified from *E. coli* cell lysate were incubated with the immobilized GST, GST-OsPRI1/2, and GST-OsTPLn in GST pull-down protein binding buffer (50 mM Tris-HCl, pH 8.0, 200 mM NaCl, 1 mM EDTA, 1%NP-40, 1 mM DTT, 10 mM MgCl_2_, 1 × protease inhibitor cocktail from Roche) for 2 h at 4°C. Proteins were eluted in the elution buffer, and the interaction was determined by western blot using anti-His antibody and anti-GST antibody (TransGen Biotech).

### Transient expression assays

The nGFP driven by a synthetic promoter which consists of five repeats of GAL4 binding motif and the minimal CaMV 35S promoter was described previously (Li et al., 2022). The GAL4 DNA binding domain was fused with mCherry containing a nuclear localization signal to generate 35S:BD-nmCherry. OsPRI1 was fused with BD-nmCherry as the effector. The promoter of OsIRO2 was used to drive nGFP as a reporter.

The promoter of *OsIRO2* was used to drive nGFP as a reporter. MYC-OsIRO3(m) and MYC-OsPRI1 were respectively cloned downstream of the 35S promoter to generate 35S:MYC-OsIRO3(m) and 35S:MYC-OsPRI1 as effectors.

*Agrobacterium tumefaciens* strain EHA105 was used for plasmid transformation. Agrobacterial cells were infiltrated into leaves of *N. benthamiana* by the infiltration buffer (0.2 mM acetosyringone, 10 mM MgCl_2_, and 10 mM MES, pH 5.6). For transcription activation assay, the final optical density at 600 nm value was 1.5. Agrobacteria were mixed at the ratio as indicated and a final concentration of 0.2 mM acetosyringone was added. After infiltration, plants were placed in the dark at 24°C for 48 h before fluorescence observation and RNA extraction. The transcript abundance of *GFP* was normalized to *NPTII*.

### Western blot

For total protein extraction, samples were ground to a fine powder in liquid nitrogen and then resuspended and extracted in protein extraction buffer (50 mM Tris, 150 mM NaCl, 1% NP-40, 0.5% sodium deoxycholate, 0.1% SDS, 1 mM PMSF, 1 x protease inhibitor cocktail [pH 8.0]). Sample was loaded onto 12% SDS-PAGE gels and transferred to nitrocellulose membranes. The membrane was blocked with TBST (10 mM Tris-Cl, 150 mM NaCl, and 0.05% Tween 20, pH8.0) containing 5% nonfat milk (TBSTM) at room temperature for 60 min and incubated with primary antibody in TBSTM (overnight at 4°C). Membranes were washed with TBST (three times for 5 min each) and then incubated with the appropriate horseradish peroxidase-conjugated secondary antibodies in TBSTM at room temperature for 1.5 h. After washing three times, bound antibodies were visualized with ECL substrate.

## ACKNOWLEDGMENTS

We thank the Institutional Center for Shared Technologies and Facilities of Xishuangbanna Tropical Botanical Garden, CAS for assistance in the determination of metal contents. We also thank the Crops Conservation and Breeding Base of XTBG for rice planting.

## Finding

This work was supported by the National Natural Science Foundation of China (32001533).

## AUTHOR CONTRIBUTIONS

G.L. conceived the project. C.L. and Y.L. constructed plasmids. C.L. characterized plants, determined gene and protein expression, and conducted cellular assays. C.L., Y.L. and P.X. grew rice and analyzed data. C.L and G.L. wrote the manuscript. All authors discussed and approved the manuscript.

## SUPPORTING INFORMATION

Additional supporting information may be found online in the Supporting Information section at the end of the article.

**Supplemental Figure S1.** OsIRO3 does not bind to the promoter of *OsFIT*.

**Supplemental Figure S2.** Expression of *OsIRO3*(*m*) in the overexpression plants.

**Supplemental Table S1.** Primers used in this paper.

