## Supplementaf information for "OsIRO3 negatively regulates Fe homeostasis by repressing the expression of *OsIRO2*"

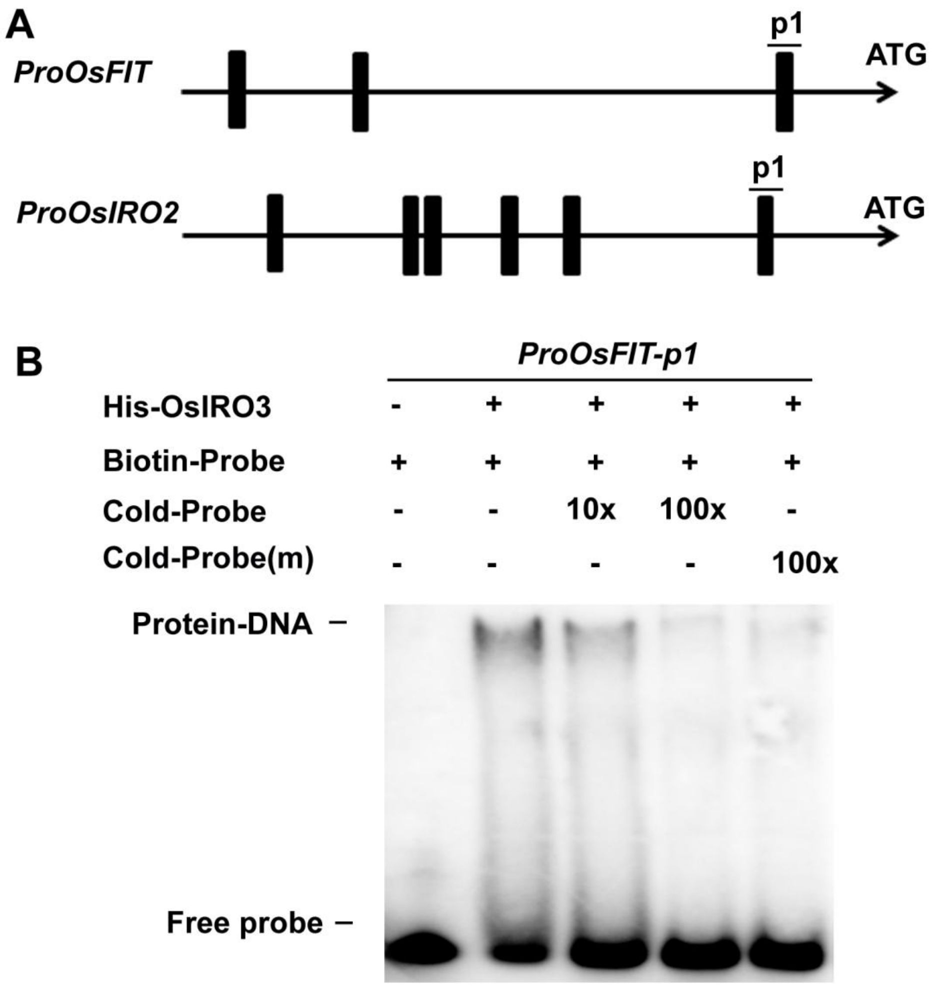


**Supplemental Figure S1.** OsIRO3 does not bind to the promoter of *OsFIT*.

(A) E-boxes in the promoter. 1kbp sequences upstream of the translation start site of OsFIT and OsIRO2 were shown. The black bar indicates E-box.

(B) EMSA assays. Biotin-labeled DNA probe was incubated with the recombinant His-OsIRO3 protein. An excess of unlabeled probe (Cold-Probe) or unlabeled mutated probe (Cold-Probe-m) was added to compete with labeled probe (Biotin-Probe). Biotin-probe incubated with His protein served as the negative control.

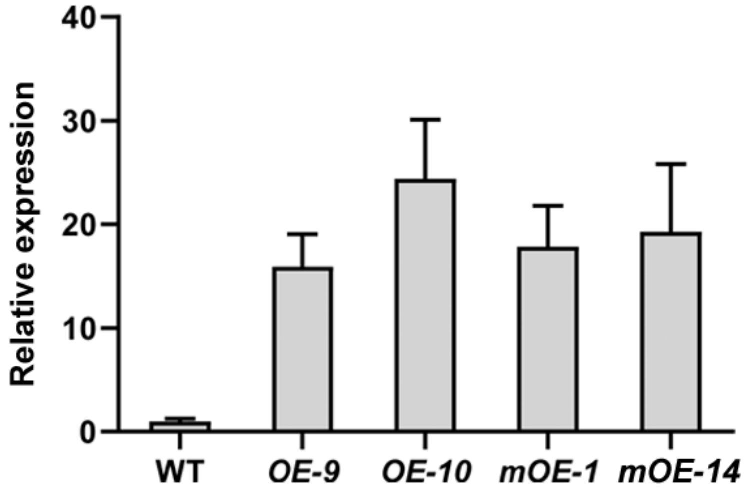


**Supplemental Figure S2.** Expression of *OsIRO3*(m) in the overexpression plants.

Shoot of ten-day-old seedlings grown in +Fe solution were used for RNA extraction and qRT-PCR. The numbers above the bars indicate the corresponding mean values. Error bars represent the SD ($n$ = 3).

**Supplemental Table S1.** Primers used in this paper.

| Name | Sequence | Template | Target vector |
|---|---|---|---|
| **For overexpression plants** | | | |
| HA-BamHI-F | TTTggatccATGGAGTACCCATACGACGTACCA | pGAD-OsIRO3(m) | pUN1301-OsIRO3(m) |
| OsIRO3-SacI-R | AAAgagctcCTACATTTGCTCATCTTCCATTCTTGGAAG | | |
| **For yeast-2-hybrid** | | | |
| GBK-OsIRO3-F | AGCTGATCTCAGAGGAGGACCTGCAtATGGTGCCGTCGGAGAGGGG | genomic DNA | pGBK-OsIRO3 |
| GBK-OsIRO3-R | GGGAATTCGGCCTCCATGGCCACTACATTTGCTCATCTTCCATTCTTGGAAG | | |
| GBK-OsIRO3n-R | GGGAATTCGGCCTCCATGGCCACGGGGAGAGGTGATCCAGT | genomic DNA | pGBK-OsIRO3n |
| GBK-OsTPLn-F | AGCTGATCTCAGAGGAGGACCTGCAtATGTCGTCGCTTAGCAGGGAGCT | genomic DNA | pGBK-OsTPLn |
| GBK-OsTPLn-R | CCCGGGAATTCGGCCTCCATGGCCACGCATTTGTCATCCAACCCGC | | |
| GBK-OsTPR1n-F | AGCTGATCTCAGAGGAGGACCTGCAtATGTCGTCGCTCAGCCGGGAGCTC | genomic DNA | pGBK-TRP1n |
| GBK-OsTPR1n-R | CCCGGGAATTCGGCCTCCATGGCCAGCAGCATTTGCCATCCAACCCGCT | | |
| GBK-OsTPR2n-F | AGCTGATCTCAGAGGAGGACCTGCAtATGTCGTCGCTGAGCCGGGAG | genomic DNA | pGBK-TRP2n |
| GBK-OsTPR2n-R | CCCGGGAATTCGGCCTCCATGGCCATCCAGCCAGCCAGAGGTGGGA | | |
| GAD-OsIRO3-F | CATACGACGTACCAGATTACGCTCAtATGGTGCCGTCGGAGAGGGG | genomic DNA | pGAD-OsIRO3 |
| GAD-OsIRO3-R | TGGAATTCACTGGCCTCCATGGCCACTACATTTGCTCATCTTCCATTCTTGGAAG | | |
| GAD-OsIRO2-F | CATACGACGTACCAGATTACGCTCAtATGGAGCAGCTGTTCGTCGACGA | genomic DNA | pGAD-OsIRO2 |
| GAD-OsIRO2-R | TGGAATTCACTGGCCTCCATGGCCATTAAAGCTTTGCTTTGTTCCTGACGA | | |
| OsIRO3m-F | CACTGGAAGCGAAGGCCTTCCTAGAAGCACCC | genomic DNA | pGBK/GAD-OsIRO3m |

| | | | |
|---|---|---|---|
| OsIRO3m-R | TTCTAGGAAGGCCTTCGCTTCCAGTGGCTGTCTTG | | |
| **For transient expression assays** | | | |
| ProIRO2-F | cgactctagaggatccccgggcgACACCCTTAGACCTTATATTTTACCG | genomic DNA | p28-ProIRO2-nGFP |
| ProIRO2-R | gaattaattccgctttatccatGCTTGCTTGTCGCTAGAGAGA | | |
| p30-MYC-BamHI-F | CTTTCGCGAGCTCGGTACCCGGGATGGAGGAGCAGAAGCTGATCT | pGBK-OsIRO3(m) | p30-MYC-OsIRO3(m) |
| p30-BD-R | CATGCCTGCAGGTCGACTCTAGAGTTATGCTAGTTATGCGGCC | | |
| p30-MYC-BamHI-F | CTTTCGCGAGCTCGGTACCCGGGATGGAGGAGCAGAAGCTGATCT | pGBK-OsPRI1 | p30-MYC-OsPRI1 |
| p30-BD-R | CATGCCTGCAGGTCGACTCTAGAGTTATGCTAGTTATGCGGCC | | |
| pTG10-MYC-F | GTTGGGTCTGGCGGTGGCTCCGAGGAGCAGAAGCTGATCTCAG | pGBK-OsIRO3 | pTG10-OsIRO3 |
| pTG10-GBK-R | GGAGGCCTGGATCGACTAGTCGGTTATGCTAGTTATGCGGCC | | |
| pTG10-MYC-F | GTTGGGTCTGGCGGTGGCTCCGAGGAGCAGAAGCTGATCTCAG | pGBK-TPLn | pTG10-OsTPL |
| pTG10-GBK-R | GGAGGCCTGGATCGACTAGTCGGTTATGCTAGTTATGCGGCC | | |
| pTG11-HA-F | gctgaagctagtcgactctagccATGGAGTACCCATACGACGTAC | pGAD-OsIRO3 | pTG11-OsIRO3 |
| pTG11-OsIRO3 | tccgccaccagaccctccaccCATTTGCTCATCTTCCATTCTT | | |
| pTG11-HA-F | gctgaagctagtcgactctagccATGGAGTACCCATACGACGTAC | pGAD-OsPRI1 | pTG11-OsPRI1 |
| pTG11-OsPRI1 | tccgccaccagaccctccaccCGCGACAGGCGGGCACGCTTC | | |
| pTG11-HA-F | gctgaagctagtcgactctagccATGGAGTACCCATACGACGTAC | pGAD-OsPRI2 | pTG11-OsPRI2 |
| pTG11-OsPRI2 | tccgccaccagaccctccaccTGCAACCGGCGGCCACATCAC | | |
| **For qRT-PCR** | | | |
| q-OsIRO2-F | GAAGGTCTTCACTTCATCAGTTCA | | |
| q-OsIRO2-R | TGATCGTTCCTTCACTTCTCTG | | |
| q-OsFIT-F | TGGAGTCGCTGTCGTGCTTCACC | | |
| q-OsFIT-R | TCAGGAGATCTGGACCGTCGGT | | |
| q-OsNAS1-F | GTGGTTCTGCCGGTGGTC | | |
| q-OsNAS1-R | AGACGGACAGCTCCTTGTTG | | |
| q-OsNAS2-F | CGTCTGAGTGCGTGCATAGT | | |
| q-OsNAS2-R | CACAAACACAAACCGATACCA | | |

| | |
|---|---|
| q-OsYSL15-F | ATCTCACCTTGACATCGCCG |
| q-OsYSL15-R | AAACGCCCTGTAGAACAGCA |
| q-OsDMAS1-F | AATCCAAGGGCAAGACCGTAG |
| q-OsDMAS1-R | CTTCACGATCAGGCAGTCCC |
| q-OsNAAT1-F | AGACCAGGCTACCCAAACTATG |
| q-OsNAAT1-R | CACCTCTTTGATAGCGATCC |
| q-OsTOM1-F | AGAGGTGCTGCAAATGGCATAT |
| q-OsTOM1-R | ATCATTTGATCCCCTGGGAAGA |
| q-OsIRT1-F | TTCGCCGTCGTCAAGGC |
| q-OsIRT1-R | GGCGAGGTGAGGTTGTTGA |
| q-OsACTIN1-F | ACACCGGTGTCATGGTCGG |
| q-OsACTIN1-R | ACACGGAGCTCGTTGTAGAA |
| q-OsOBP-F | GGCGCCGAAGAAGCTATTG |
| q-OsOBP-R | GTTGCTCTTCAAGCCTGCTC |
| **For EMSA** | |
| pOsFIT-F | TCAAGTAGTATTAGTCAATTGGTC ATCACATTTCG |
| pOsFIT-R | CGAAATGTGATGACCAATTGACT AATACTACTTGA |
| mpOsFIT-F | TCAAGTAGTATTAGTGGCTTGGT CATCACATTTCG |
| mpOsFIT-R | CGAAATGTGATGACCAAGCCACT AATACTACTTGA |
| pOsIRO2-F | CCTCCCTAGCTTGGCAGAATAGT TAATGTTAATCACCTGATTAAA |
| pOsIRO2-R | TTTAATCAGGTGATTAACATTAAC TATTCTGCCAAGCTAGGGAGG |
| mpOsIRO2-F | CCTCCCTAGCTTGGCAGAATAGT TAATGTTAATGTCCTGATTAAA |
| mpOsIRO2-R | TTTAATCAGGACATTAACATTAAC TATTCTGCCAAGCTAGGGAGG |
